# The role of nature conservation and commercial farming in psychological distress among rural Ugandans

**DOI:** 10.1101/2021.06.08.446718

**Authors:** Thomas Pienkowski, Aidan Keane, Eugene Kinyanda, Birthe Loa Knizek, Caroline Asiimwe, Geoffrey Muhanguzi, E.J. Milner-Gulland

## Abstract

Mental illness is a leading contributor to the global burden of disease, but there is limited understanding of how it is influenced by socio-ecological context, particularly in the global south. We asked how interactions with ecological systems influence stressors associated with psychological distress in a rural Ugandan case study. We conducted and thematically analyzed 45 semi-structured interviews with residents of Nyabyeya Parish, Masindi District. Our results suggest that poverty and food insecurity were the primary reported causes of *“thinking too much”* and related idioms of psychological distress. The expansion of commercial agriculture may have been associated with the contraction of subsistence farming, reportedly exacerbating poverty and food insecurity among poorer households but contributing incomes to wealthier ones. Furthermore, households bordering a conservation area reported that crop losses from wildlife contributed to food insecurity. However, forest resources were important safety nets for those facing poverty and food insecurity. Our study suggests how two globally prevalent land uses – commercial agriculture and nature conservation – may influence social determinants of psychological distress in our study area. Psychological distress does not necessarily imply mental disorder. Nonetheless, exploring socially-mediated interactions with ecosystems may help explain the etiology of psychological distress. Furthermore, we suggest opportunities to manage socio-ecological systems to support mental health, such as promoting equitable access and control of livelihood resources. We also highlight co-benefits and trade-offs between global sustainability goals that could be managed for mental health, and why these should be recognized in the anticipated ‘New Deal for Nature.’

**Highlights:**

- Food insecurity and poverty were major stressors reportedly causing psychological distress, characterized as *“thinking too much.”*
- Complex interactions between conservation, commercial agriculture, and poverty influenced psychological distress.
- Commercial agriculture displaced subsistence farming, exacerbating poverty and food security among poorer households.
- Conserved forests were both home to crop-raiding wildlife and sources of income and food, affecting poverty and food insecurity.
- Mental health needs to be included in assessments of the role of the ecosystems in public health.

## 1. Introduction

Global environmental change poses new and significant threats to human health (Myers, 2017). These threats are diverse, from upstream human activity increasing diarrhea risk in the forested tropics to climate change reducing the nutritional value of crops globally (Herrera et al., 2017; Myers et al., 2014; Pienkowski et al., 2017). These threats to health are also expected to grow in the face of worsening biodiversity loss, ecological degradation, and climate change (Whitmee et al., 2015). Consequently, health now features in mainstream environmental discourse. For example, the sixth UN’s flagship *Global Environment Outlook* report states, *“a healthy planet is a necessary foundation for human physical, psychological, social, economic and emotional health and well-being”* (UNEP, 2019a, p. 5). Accordingly, many now claim that environmental protection is necessary to meet health-related Sustainable Development Goals (e.g., Myers, 2017; Wood et al., 2018).

Health is defined as a state of physical, mental, and social well-being, with illness being the disruption of health (Boyd, 2000). Mental illness is a core determinant of health, being among the largest cause of years lived with disability, mostly attributed to common mental disorders such as depression (NCCMH, 2011; Vigo et al., 2016). Despite its associated large and growing burden of disease, mental illness continues to be neglected within public health agendas. For instance, mental, neurological, and substance use disorders and self-harm accounted for 12% of disability-adjusted life years, and 32.4% of years lived with disability globally in 2015, but only 2% of public health spending (Vigo et al., 2019). Mental health is particularly neglected in low-income countries, where mental disorders are the largest cause of years lived with disability, but less than 5% of those in need receive adequate care (IHME, 2018; Vigo et al., 2019).

This neglect has been called a *“failure of humanity*” and appears to be reflected in research on the health impacts of environmental change (Kleinman, 2009, p. 603). For example, among 1,306 articles mentioning natural environments and human health, illness, or diseases, only 5% included terms associated with mental health or common mental disorders in their title or abstract (see Supplementary Information 1: Article search method, NCCMH, 2011; 2011; Scopus, 2020). A growing body of research explores the psychological impacts of climate change and air, soil, and water pollution (Cianconi et al., 2020; Frumkin & Haines, 2019; Middleton, Cunsolo, Jones-Bitton, Wright, et al., 2020). However, relatively few studies examine the role of habitats, biodiversity, and other ecological factors, which we henceforth refer to as ecosystems. Much of the research connecting ecosystems and mental health has narrowly focused on the psychological benefits of green and blue spaces (e.g., Bratman et al., 2019). We suggest two limitations with this literature; the neglect of the rural global south and a limited understanding of the indirect ways that ecosystem change influences social determinants of mental illness.

First, much of the evidence linking ecosystems and mental illness comes from the global north, with some important exceptions (e.g., Tomita et al. (2017), which explores green environments and depression incidence in South Africa). Many of these reported links are desirable, such as the benefits of urban green spaces for New Zealanders (Nutsford et al., 2013). Nevertheless, this evidence largely overlooks the experience of nearly three billion people living in the rural global south (World Bank, 2020). These experiences can differ from those typically reported in the global north. For instance, studies in India describe how living close to large mammals can lead to traumatic encounters potentially detrimental to mental health (Barua et al., 2013; Chowdhurym et al., 2016; Jadhav & Barua, 2012). These experiences vary because of differences in both the social as well as the ecological context (Lawrence et al., 2019). For instance, the traumatic encounters mentioned above occur because of the proximity between conservation areas and rural residents (e.g., Chowdhurym et al., 2016). This point emphasizes the need to understand how mental health outcomes are co-produced through the interaction of society and ecosystems, termed ‘socio-ecological systems.’ Socio-ecological systems are linked and interdependent systems of nature and people, which are nested across scaled and evolve over time (Bouamrane et al., 2016; Liu et al., 2007). A vast body of research explores how nature’s contributions – the benefits and costs that people obtain from nature – are co-produced within socio-ecological systems (IPBES, 2020). For instance, agriculture, including both subsistence and industrialized farming, involves closely coupled human and natural systems that underpin human well-being (TEEB, 2018). Consequently, a socio-ecological systems perspective may help understand how and why relationships between ecosystems and mental health may differ between places.

Second, within the health literature, people’s social, economic, and built environmental contexts are all considered ‘social determinants’ of mental illness (Patel et al., 2018). Much of the evidence connecting ecosystems to mental health focuses on *direct* relationships, such as the benefits of urban green spaces (Hartig et al., 2014). However, many of nature’s contributions are likely to influence these social determinants of mental illness. In other words, nature contributions may *indirectly* alter the risk of mental illness through their effects on people’s social, economic and cultural context. However, it appears that little empirical research explores socially-mediated relationships between ecosystems and mental illness. In contrast, growing climate change research explicitly examines the role of social determinants of mental illness (e.g., Berry et al., 2018). For instance, one review identified multiple mechanisms by which drought could indirectly affect mental illness, such as financial stress exacerbating family tensions, triggering depression (Vins et al., 2015). Another study suggested that abnormal weather patterns influenced Inuit identity, social relationships, and culture in ways harmful to their mental well-being (Middleton, Cunsolo, Jones-Bitton, Shiwak, et al., 2020). There are likely to be myriad diverse, widespread, and important links between ecosystems and mental illness that have been overlooked. For instance, soil biodiversity loss may compromise agricultural yields, thereby triggering food insecurity, a known social determinant of mental illness (Jones, 2017; Lund et al., 2018; Weaver & Hadley, 2009). These connections might be particularly important in parts of the rural global south where nature’s contributions to people can be most valuable.

Taken together, these limitations within the literature have produced an incomplete and geographically-biased understanding of the ways that interacting with ecosystems might influence mental health and illness. Accounting for the role of ecosystems may help explain variation in mental illness between groups, such as the observed but unexplained differences between communities in Uganda (Kinyanda et al., 2013; Kinyanda et al., 2017; Kinyanda, Woodburn, et al., 2011). Furthermore, understanding how interacting with ecosystems may influence mental health might also help predict and avoid negative impacts of ecological degradation. A more nuanced and locally appropriate understanding of these interactions may reveal new opportunities to jointly address global ecological and mental health priorities (Patel et al., 2018; Whitmee et al., 2015).

This paper aims to demonstrate the value in such an holistic approach. It illustrates previously unexplored ways in which interacting with ecosystems may influence psychological distress – a state of non-specific emotional disturbance – through a rural Ugandan case study. The case study is conducted among nine rural communities in Nyabyeya Parish, in western Uganda. This Parish borders Budongo Forest Reserve, a valuable site for Ugandan nature conservation, but which has experienced extensive recent land-use change. Therefore, the relationship between ecosystems and people’s experiences of psychological distress may be particularly acute. Within the case study, we ask:

1. How do Nyabyeya’s residents describe their experiences of distress?
2. What are the perceived stressors causing this distress among residents?
3. What are the perceived roles of interactions with ecosystems, and the socio-ecological setting, in producing these stressors?

### 1.1. Conceptual framework

We developed a conceptual framework, describing how interactions between social systems and ecosystems might influence social determinants of psychological distress, to guide the semi-structured interviews within the case study. The conceptual framework has three primary components related to; a) how stressors may elevate the risk of psychological distress, b) the types of stressors people experience, c) the ways people’s socio-ecological context can influence these stressors (see Supplementary Information 2: Conceptual framework).

The first component of the framework describes how excessive exposure to stressors may increase the risk of psychological distress (Figure 1). Stressors are defined as events, conditions, or forces that result in emotional or physical stress (VandenBos, 2007). Psychological distress has been defined as a state of emotional disturbance that impairs day-to-day activities and social functioning (Drapeau et al., 2012). A stage-based model of mental illness suggests a spectrum from mental health, through diffuse and non-specific psychological distress, to definable syndromes (McGorry et al., 2014; McGorry & van Os, 2013). At the early stages, individuals may experience a wide range of non-specific symptoms, such as fatigue, anxiety, or insomnia. At later stages, these can become more severe, specific, and resistant to treatment, eventually passing diagnostic thresholds for mental illness (Patel et al., 2018). Individuals can move in both directions along this spectrum, and experiences of stressors over an individual’s life-course can be risk factors within this stage-based model (Johnstone et al., 2018; Lund et al., 2018). Furthermore, the pathogenic effect of these stressors, and an individual’s predisposition to mental illness, is partly a function of their genetic, neurodevelopmental, and psychological characteristics (Goh & Agius, 2010; McGorry et al., 2014).

**Figure 1.**
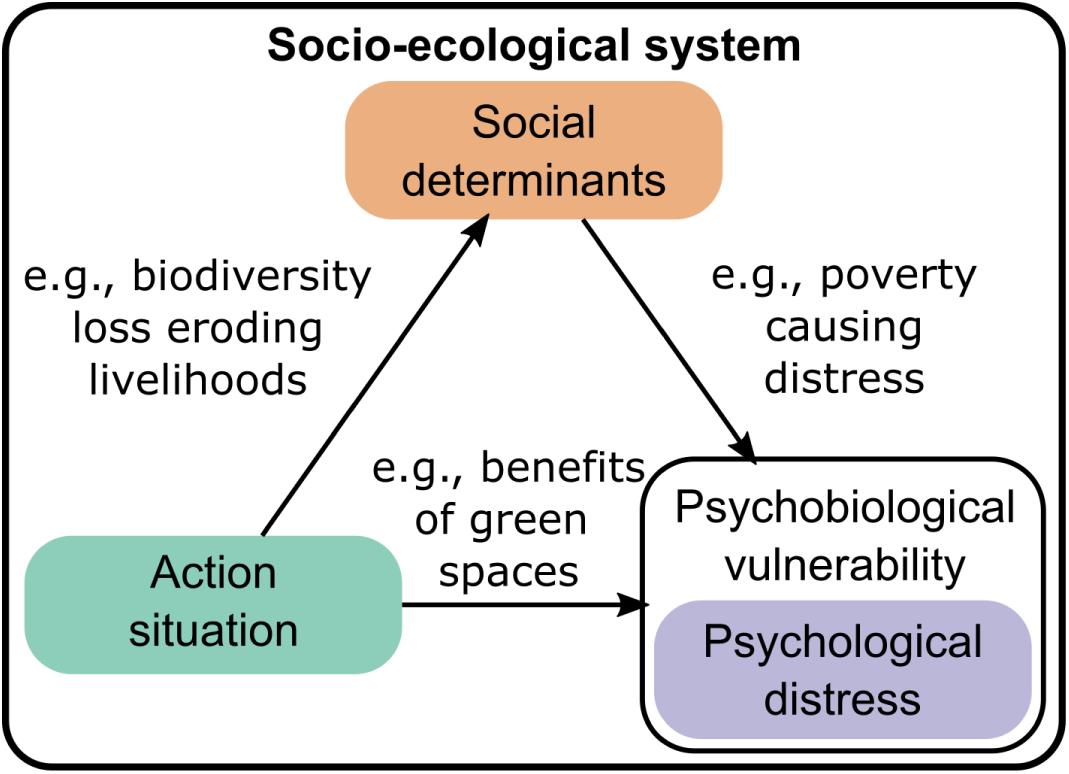
The direct and indirect ways that interacting with ecosystems could influence the risk of psychological distress, depending on an individual’s psychobiological characteristics, within a socio-ecological system. The socio-ecological system includes multiple interacting elements; those of interest to a researcher, decision-maker, or other user are termed ‘action situations.’

As an exploratory study using qualitative methods, we seek to identify stressors that individuals associate with non-specific psychological distress. We emphasize that further research is required to understand if these translate into greater risk of later-stages syndromes. We explored experiences of distress using locally appropriate terms. These idioms of distress are culturally and socially situated ways of experiencing and describing distress (Nichter, 2010).

The second component describes broad categories of stressors faced by populations experiencing poverty, drawing on the Voices of the Poor initiative (Narayan et al., 2000). This initiative identified five categories of stressor: *material lack and want*; *physical ill-being*; *bad social relations*; *insecurity and vulnerability*; and *powerlessness, frustration, and anger*. These broad categories represent potential social determinants of psychological distress among those experiencing poverty.

The final component describes how the interaction of social and ecological systems defines the context of people’s lives, including the stressors they face. Ostrom and colleagues provided a framework for organizing and structuring the many features found in socio-ecological systems within a multi-level framework (Colding & Barthel, 2019; Ostrom, 2007). Social elements of this framework include governance systems and actors, and their socioeconomic, cultural, religious, and other characteristics. Ecological aspects of this framework include natural resource systems and units and their characteristics. Interactions of these elements over time result in the co-evolution of socio-ecological systems (Liu et al., 2007). While providing an overview of the interconnected elements of a system, this framework is often used to examine a specific phenomenon of interest within an ‘action situation.’ Within our study, the action situation was not known in advance, and so is not specified at this stage. We use this framework as a tool to think systematically about how social and ecological variables interact within an action situation to affect the stressors people face. As such, the first and second components of the framework are nested within the socio-ecological system component. In summary, our framework represents how interactions with ecosystems may influence social determinants of psychological distress, as expressed in locally appropriate terms.

## 2. Methods

### 2.1. Study site: Nyabyeya Parish and the surrounding areas

The scope of the case study includes Nyabyeya Parish, the nine communities within it, and the surrounding area (Figure 2). This site was chosen because many residents are subsistence farmers or make a living through forest-resource use, where ecological processes can be instrumental to livelihood and well-being outcomes. However, these systems are also changing as the result of commercial agricultural expansion and forest cover loss with possible implications for residents’ well-being.

**Figure 2.**
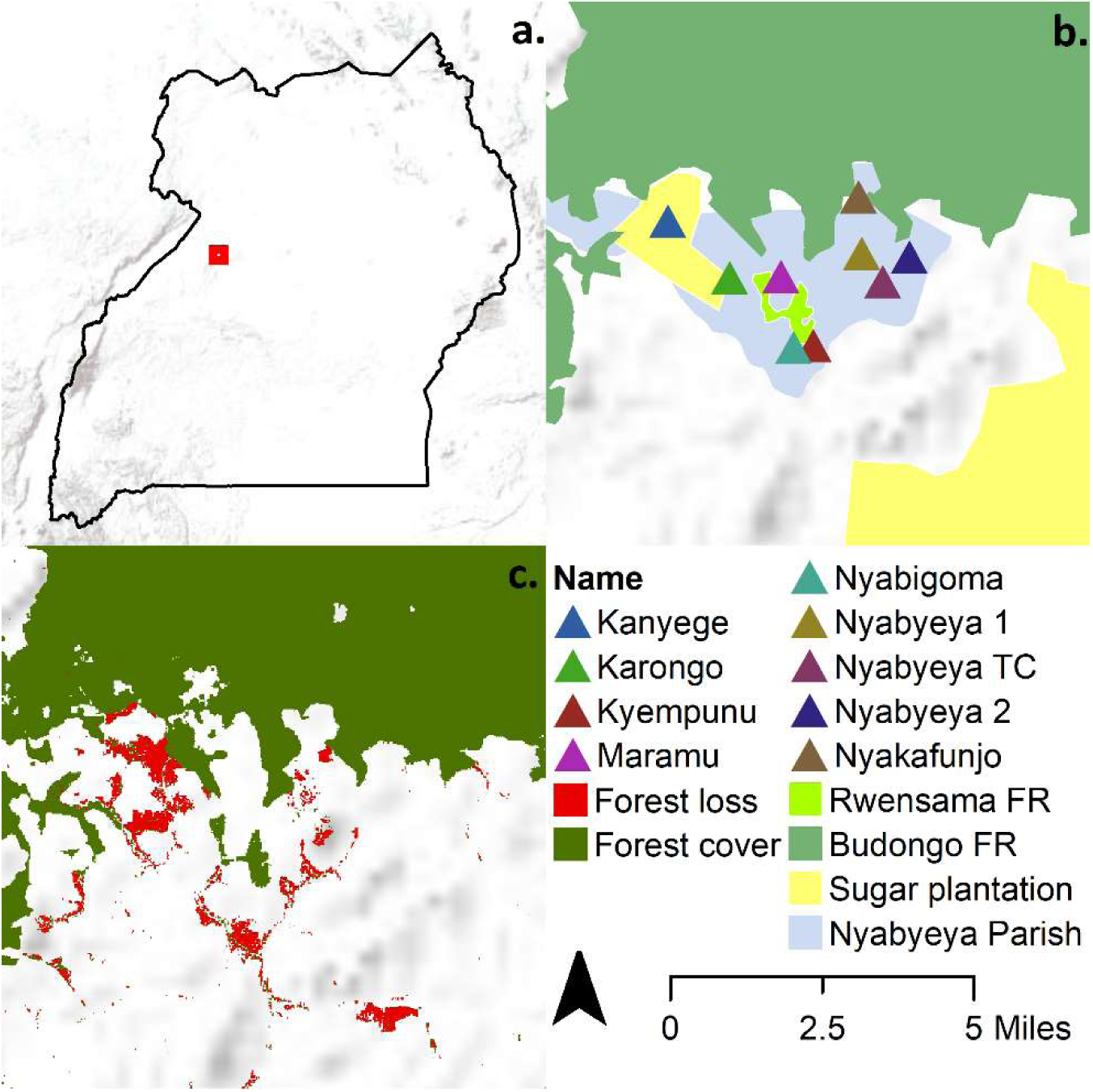
Panel a. describes the location of the study site within Uganda (Esri et al., 2009). Panel b. describes the study area, which includes Nyabyeya Parish and its nine communities, the Budongo and Rwensama Forest Reserves (FR), and the indicative location of large-scale commercial sugarcane estates (GADM, 2018; adapted from Kyongera, 2015; UNEP-WCMC & IUCN, 2020). Panel c. describes forest loss between 2000 and 2016 and forest cover (>75% tree cover) in 2016 (adapted from Hansen et al., 2013).

Budongo Forest Reserve, neighboring the smaller and fragmented Rwensama Forest Reserve, was gazetted for timber production by the British in the 1930s (Paterson, 1991). By 1960 Budongo Sawmills was the largest timber producer in the country and was a driver of in-migration before closing in the late 1990s (Babweteera et al., 2018; Babweteera et al., 2012). The refurbishment of Kinyara Sugar Works in the 1990s was another pull factor for migrants and now represents an important source of employment in mid-western Uganda (Babweteera et al., 2012). As a result, Nyabyeya Parish is linguistically and culturally diverse while being home to the indigenous Banyoro people.

Since 2000, the Ugandan government has adopted policies seeking to alleviate rural poverty through agricultural industrialization (Nabwire, 2015). These policies, accompanied by increased global demand, contributed to the expansion of commercial sugarcane farming in Masindi District and other parts of Uganda (Jeary et al., 2018). Most of Nyabyeya’s residents are small-scale farmers who grow food crops for subsistence and sale (Babweteera et al., 2018). However, some wealthier land-owning residents have transitioned to small-scale contract sugarcane farming (also referred to as ‘out-growers’) to supply Kinyara Sugar Works, which supplies national and international markets (Twongyirwe et al., 2017). Nyabyeya also includes several large-scale commercial sugarcane estates that also supply Kinyara Sugar Works. Nyabyeya Parish has experienced significant forest loss over the last 50 years, much of which occurred before the 2000s and outside the boundaries of the forest reserves (Twongyirwe et al., 2017). As a result, the forest edge now largely coincides with the boundaries of the two reserves. Budongo Forest Reserve forms an important component of the Albertine Rift, a valuable region for conservation in Africa (Plumptre et al., 2007). However, the smaller Rwensama Forest Reserve appears to be heavily degraded. Moreover, illegal timber extraction, charcoal production, and hunting occur within both forests. The latter often involves the use of snares, which are a significant threat to endangered chimpanzees and other wildlife (Babweteera et al., 2018). Nevertheless, Nyabyeya’s forests are important sources of firewood, medicine, wild food, and other products (Babweteera et al., 2018; Eilu & Bukenya-Ziraba, 2004; Tumusiime et al., 2010). At the same time, chimpanzees, baboons, and other wildlife are reportedly significant causes of crop losses for farmers on the edges of the two forest reserves (Hill, 2015).

There appear to be no nationally representative measures of levels of psychological distress in Uganda. However, one regional study suggested that 0.9% of 6,663 respondents’ in southwestern Uganda screened positive for severe psychological distress (Kinyanda, Waswa, et al., 2011). More broadly, depressive disorders were the third-largest source of years lived with disability in 2016 (IHME, 2018). One study of 4,660 Ugandans estimated that 29.3% of respondents might have met the diagnostic criteria for current major depressive disorder in 2003 or 2004 (Kinyanda, Woodburn, et al., 2011). Furthermore, formal care for those with poor mental health is limited (Molodynski et al., 2017). The closest mental healthcare provider to Nyabyeya Parish is Hoima Regional Referral Hospital, over an hour’s drive away. Nevertheless, we expect Nyabyeya residents’ mental health to be typical of that found in other rural Ugandan populations.

### 2.2. Ethical considerations

Ethical approval was granted by the Uganda National Council of Science and Technology (Ref. SS6007) of the Government of Uganda, and the Central University Research Ethics Committee (Ref. R63458) at the University of Oxford (see Supplementary Information 3: Ethical considerations).

### 2.3. Study population and sample

The study population includes the primary decision-making male and female family members over the age of 18 in the nine communities in Nyabyeya Parish in western Uganda. These individuals were selected because they were expected to feel responsible for other household members, and thus distressed by stressors affecting their family. Furthermore, these individuals were also considered knowledgeable about household and community conditions and were a relatively easy group to identify and sample consistently. This sample includes indigenous Banyoro and in-migrant subsistence and small-scale contract sugarcane farmers, non-farmers, and landless commercial agricultural workers.

A total of 45 semi-structured interviews, lasting 118 minutes each on average, were conducted in the nine communities from September to November 2019. Respondents were purposively sampled by walking through each community and observing households. This purposeful sampling of respondents sought to capture variation in gender, age, socioeconomic status, and forest proximity. The 45 interviews were deemed sufficient to capture much of the variation in these key characteristics within Nyabyeya Parish. Moreover, we prioritized conducting fewer in-depth interviews over numerous superficial ones. The target sample was five interviews in each of the nine communities. One interview was excluded from Kanyege community at the request of the respondent. An additional interview was conducted in Nyabigoma community since another interview in this community was brief and covered only some of the interview guide themes (see Supplementary Information 4: Sampling effort).

### 2.4. Data collected

Data were collected through semi-structured interviews, following an interview guide (see Supplementary Information 5: Interview guide). This approach provided the opportunity for new themes to emerge while ensuring consistency between interviews. The interview guide included broad themes from the conceptual framework in three sections. The first section discussed specific themes derived from the socio-ecological system framework, relating to actors, governance systems, natural resource systems, the broader social, economic, and political, and environmental context, and their interactions. The second section was related to the five ill-being domains found in the Voices of the Poor framework. The third section focused on the respondent’s experience of distress. During the second section, respondents were asked about perceived causes and consequences of stressors, thereby eliciting connections between the first and third sections.

### 2.5. Piloting and translation

Interviews were piloted with two individuals from a community outside the study population. The dialogue was translated between English and Kiswahili and Runyoro (the language of the indigenous Banyoro) by three research assistants during the interview, recorded on a Dictaphone. The in-situ translations were transcribed as we did not have the resources to translate the interviews verbatim, meaning that some nuances may have been lost or reinterpreted by the translator. However, the thematic analysis (discussed below) was largely semantic, focusing on what respondents explicitly stated rather than looking for latent meaning. Consequently, it is unlikely that the use of verbatim translation would have significantly changed the results.

### 2.6. Data analysis

We employed inductive thematic analysis identifying, analyzing, organizing, and reporting patterns and themes within the data (Braun & Clarke, 2006). The analysis was semantic, meaning we were interested in the surface rather than the latent meaning of what respondents said. The thematic analysis proceeded through the following broad steps (see Supplementary Information 6: Steps in the thematic analysis), conducted by the first author:

1. Familiarization with data, including through re-reading transcripts and post-scripts, and comparing word clouds between groups.
2. Generating codes and coding text through broad then specific coding in two rounds.
3. Searching and clustering into key themes, by clustering related codes into concepts.
4. Reviewing themes, ensuring consistency within themes, but discrete differences between them, and visually exploring connections between codes as networks.
5. Defining and naming themes (see Supplementary Information 7: Definition of key themes).

The expected ways that the authors’ positionality influenced the results, and steps to account for this, are discussed in Supplementary Information 8: Positionality statement.

## 3. Results

We interviewed 21 men and 24 women in a range of age groups, ethnicities, and levels of wealth (see Supplementary Information 4: Sampling effort). A broad range of themes were discussed during the interviews. Frequently occurring themes relating interactions with ecosystems to experiences of distress are presented, but other less relevant themes are not (but see Supplementary Information 9: Other pathways). In summary, experiences of *“thinking too much”* and related idioms of distress often included heart palpitations, chest and abdominal discomfort, fatigue, feeling faint, weakness, *“growing thin,”* and changes in appetite. The primary reported sources of *“thinking too much”* and related idioms were poverty and food insecurity. These were reportedly influenced by multiple factors, including changing agricultural systems and interacting with conserved forests. The following themes are the three most relevant to the research aims. First, the expansion of commercial agriculture was reportedly associated with declining subsistence farming. This decline was a perceived contributor to poverty and food insecurity among poorer households with limited land. However, engaging in commercial agriculture was a reported source of income for wealthier land-owning families, who included indigenous Banyoro and some migrants. Second, households bordering Budongo and Rwensama Forest Reserves reported crop losses from wildlife that contributed to food insecurity. These border households included those who had migrated to the area to work in Budongo Sawmills before it closed, and who appeared to settle in the highest densities around the forest edge. Finally, forest resources such as timber were also reported as sources of food and income for those experiencing poverty or food insecurity. Illegally harvested forest resources in the form of timber, animals, or charcoal appeared to be particularly important for these individuals.

### 3.1. How do Nyabyeya’s residents describe their experiences of distress?

Terms used to describe the experience of distress include *“kufikiri sana”* in Kiswahili and *“kuterageza muno”* in Runyoro, both of which translate to *“thinking too much”* in English. For instance, respondent R05 (lower-income older female) said, *“it brings me so many thoughts, I think too much.”* Other idioms of distress expressed in English include *“overthinking,” “strong thoughts,” “lots of thoughts,” “too many thoughts,”* and *“thinking a lot.”* In the following, ‘thinking too much’ refers to all of these idioms of distress used by respondents.

When asked about the experience of ‘thinking too much,’ respondents mentioned a range of symptoms (Figure 3). For instance, R29 (lower-income middle-aged male) stated, *“You find yourself growing more thin and thin. Like the way they say that too much thoughts causes pressure.”* Several reported that the experience of ‘thinking too much’ disrupted daily activities. For example, R09 (middle-income middle-aged male) said, *“You sleep now from night up till 10 am.* […] *you are supposed to wake up and get your hoe and start digging, so those are all about thoughts.”*

**Figure 3.**
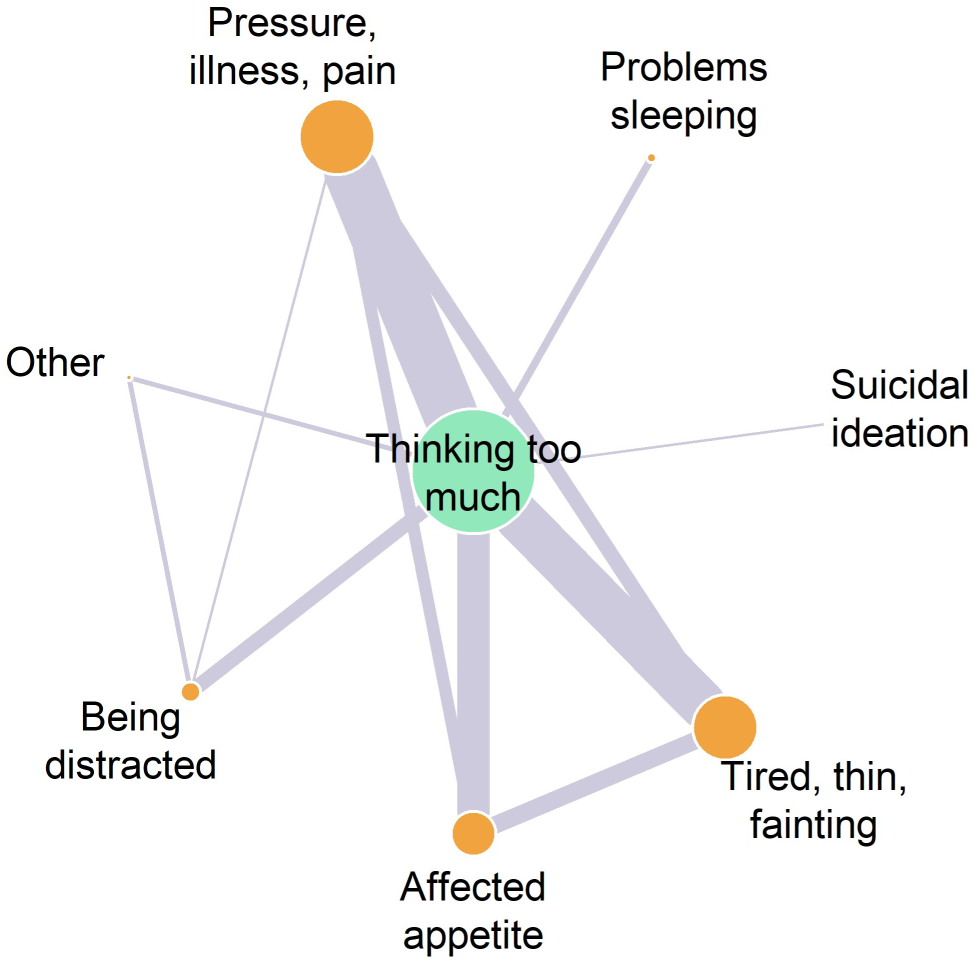
The reported experiences or symptoms of ‘thinking too much’ (and other idioms of distress). The width of the lines illustrates the relative number of interviews that reported connections between nodes. The lines represent associations reported by interviewees but do not imply directional causality. The size of the node represents the number of interviews mentioning the associated theme for that node.

Several respondents indicated that the frequency and duration of ‘thinking too much’ depended on the presence of specific stressors. For instance, when asked how to alleviate ‘thinking too much,’ R38 (middle-income older male) said, *“There is no way you can reduce those thoughts if you are still with those challenges unless those challenges are not there.”* However, several others indicated the experience of ‘thinking too much’ was more chronic, such as R27 (lower-income middle-aged female), who said, *“you cannot imagine the period that those thoughts can get finished from you.”*

### 3.2. What are the perceived stressors associated with distress among residents?

Many respondents reported that poverty, bad health, and inadequate food were associated with ‘thinking too much’ (Figure 4). For instance, when asked what the term *“overthinking”* meant, R20 (middle-income older female) responded, *“No energy for digging*, *no money brings famine, it makes you overthink. You start thinking what will I eat.”* Although being poor was used as an umbrella term for someone’s socioeconomic condition, it was often mentioned in association with not having enough money to meet essential needs. These needs included basic housing, paying for healthcare and school fees, and buying food. In the following, we therefore use the term ‘poverty’ to mean inadequate money.

**Figure 4.**
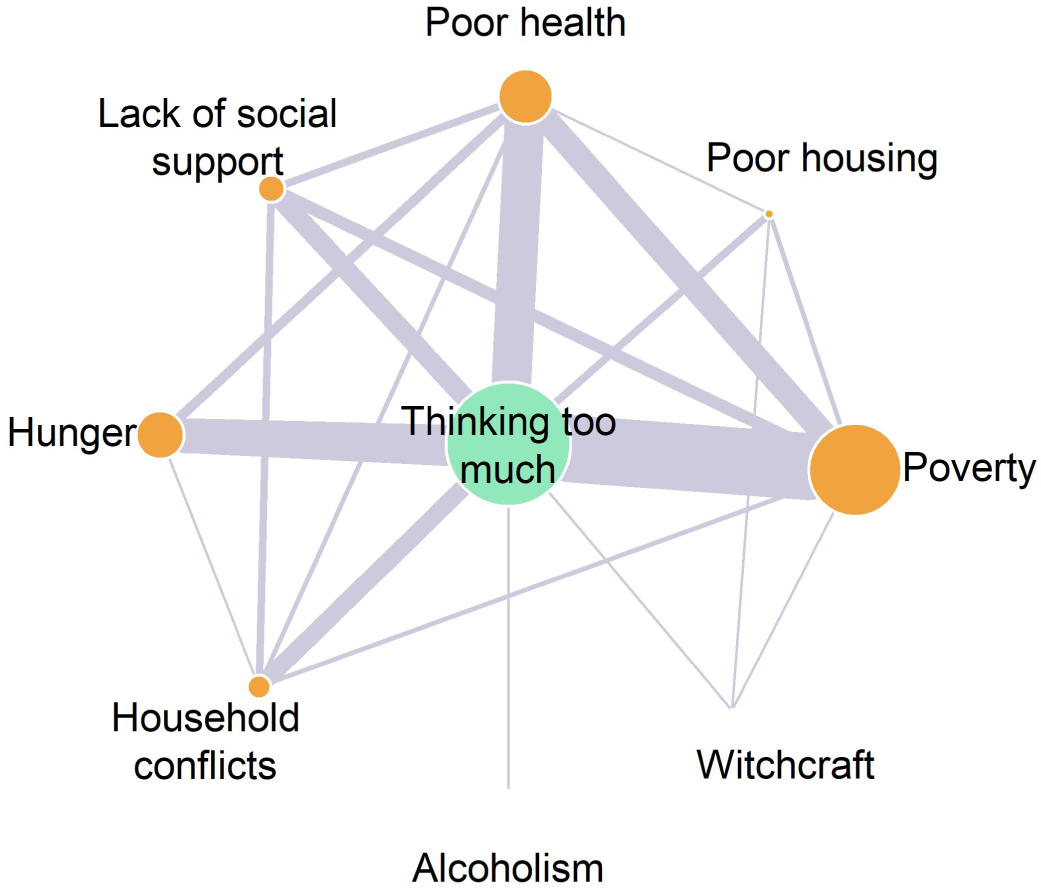
The reported sources of ‘thinking too much’ and related idioms of distress. The width of the lines illustrates the relative number of interviews that reported connections between nodes. The lines represent associations reported by interviewees but do not imply directional causality. The size of the node represents the number of interviews mentioning the associated theme for that node.

Many respondents, particularly subsistence farmers, used the terms ‘famine’ and ‘hunger’ to describe not having enough food. When asked how this affected respondents’ households, several mentioned having to reduce the size or number of meals eaten per day and eating less favored food or the same types of food in multiple meals. For instance, when asked about the experience of hunger, R13 (middle-income younger male) stated, *“You cannot eat expensive things, and if you have been eating like four cups of posho* (maize flour) *now, you end up reducing to two cups.”* Although a few respondents indicated that the current experience of hunger was a cause of ‘thinking too much,’ many more said that the prospect of food supplies running out before the next harvest season caused them distress.

### 3.3. What are the perceived roles of socio-ecological processes in these stressors?

In the following, we focus on poverty and hunger because they were the most frequently mentioned intermediaries between the respondent’s broader socio-ecological context and ‘thinking too much.’ Further evidence illustrating these results and other pathways of interest are presented in the SI (Supplementary Information 9-12).

#### 3.3.1. Poverty, farm size, and the sugarcane industry

The primary reported contributors to poverty were low farm production, poor health, and lack of employment. For instance, R29 (lower-income middle-aged male) stated, *“What has caused the poverty in this community is lack of enough land, you do not have where to dig, you just have a plot.”*

When discussing farm production, many subsistence farmers suggested there was inadequate or *“squeezed”* land. For instance, when asked to describe the history of the community, R25 (medium income middle-aged male) said, *“others are getting problems of land because it is small where they can do good farming to get good money […] that’s what is making us to suffer.”* When asked why there was inadequate land, a majority of subsistence farming respondents reported that the expansion of small-scale contract farming and large-scale commercial estates had displaced subsistence farming. For instance, R02 (middle-income middle-aged male) stated, *“you cannot struggle for a small piece of land since sugarcane has taken most parts of the land.”* Mechanisms of displacement reported by both commercial and subsistence farmers included the voluntary selling or renting of land to meet immediate needs, forced displacement by large-scale commercial estates (particularly in Karongo and Kanyege in the west), and increased prices restricting land purchase. Additionally, a large number claimed that the sugarcane industry affected land availability without explaining the process through which this happened.

However, both subsistence and small-scale contract farmers said that selling sugarcane was a source of household development. For instance, when describing the drivers of household development, R01 (middle-income middle-aged female) said, *“sugarcane growing, and after growing, they sell and get a lot of money.”* Nevertheless, many subsistence farmers reported barriers to small-scale contract farming, the most commonly mentioned of which was not having enough land. Consequently, several respondents said that the benefits of the sugarcane industry mainly accrued to wealthier households with larger amounts of land.

Several suggested that the indigenous Banyoro and early migrants owned more land and were thus more likely to be small-scale contract farmers than recent migrants. For instance, R41 (middle-income older female) stated, *“Those people that have a large amount of land, they are the people who came first and a long time ago* […] *when you look at this time, land is too expensive, and it is not easy to* [get]*.”* However, a few others also suggested that some recent and relatively wealthy migrants also purchased land for small-scale contract farming. Additionally, several respondents also reported that commercial agriculture was a pull-factor for transient migration. For instance, R33 (lower-income middle-aged female) stated, *“those people who do sugar cane cutting, they always come, and they can spend here some time but when they do close the factory, maybe for maintenance, they always go back in their places.”*

Additionally, several respondents – mostly from the communities Maramu, Nyakafunjo, and Nyabyeya 1 – described a Community Forest Management initiative. Within this initiative, some residents were permitted to grow crops and timber on National Forest Authority land. Notionally, this initiative reportedly sought to discourage illegal forest use and to provide a buffer between farms and Budongo Forest Reserve. This initiative reportedly represented a source of additional land, but its conservation effectiveness was unclear.

#### 3.3.2. Hunger, nature conservation, and the sugarcane industry

The dominant reported drivers of hunger included low farm production (linked to the sugarcane industry - as above), inadequate money to buy food, and crop losses. Many subsistence farmers said that unexpected and unseasonal rains in June and July 2019 left beans, maize, and other crops rotting in their fields. Many also reported that crop-raiding by wildlife (baboons, chimpanzees, and wild pigs) caused crop losses. For instance, R16 (lower-income younger female) stated that *“Wild animals are good at spoiling most of our food crops.”* Both reserve-adjacent and non-reserve-adjacent interviewees said that those at the forest edge were most affected by crop-raiding.

In response to crop-raiding, many subsistence farmers reported having to guard their crops, which several respondents said disrupted other income-generating activities or leisure time. Consequently, several subsistence farmers also indicated negative attitudes towards wildlife and a desire to trap or kill wild animals. However, it was unclear if respondents acted upon these desires. For instance, R18 (middle-income middle-aged female) stated, *“*[wild animals] *cannot leave you to eat your food, and you are not allowed to kill them, meaning you just suffer. But if you get time, you can go and wait for them. You chase them.”* Similarly, when asked about setting traps in the forest, R03 (lower-income middle-aged male) said, *“because if it is disturbing your crops, the best thing is to kill it, you eat, and even your crops get saved.”*

#### 3.3.3. The importance of the forest during hunger and poverty

As well as the threats posed by crop-raiding wildlife, many said that some subsistence farmers and landless young men used the forest to help cope with hunger or poverty. This forest use included legal activities such as harvesting forest foods, and illegal activities such as burning charcoal, hunting, and timber harvesting. Several respondents reported that timber harvesting was often conducted by landless young men, paid by wealthier households. Similarly, several respondents said that charcoal was typically produced by poorer households and sold to wealthier ones. For instance, R02 (middle-income middle-aged male) said, “[people are struggling to survive] *as the sugarcane is the most* [common] *crop grown on the ground, so* [food crop] *gardens are few.”* R02 then proceeded to state, *“and that is why people are entering in the forest just stealing the forest, just burning charcoal, collecting firewood in days that are not allowed.”* Hunting and harvesting of forest foods reportedly directly contributed to diets. Furthermore, income generated from forest product harvesting was reportedly used to purchase food and meet other needs. However, many respondents also mentioned the range of risks associated with illegal forest use, including corporal punishment, fines, and imprisonment. For example, R33 (lower-income middle-aged female) said, *“If you are caught, the only thing is caning and getting imprisoned and you getting out of prison, meaning that you are going to pay some money.”* However, no respondents explicitly identified these risks as stressors associated with ‘thinking too much.’

Additionally, interviewees were asked if the extent of forest cover around communities had changed since they first arrived in the area or were children. Some older respondents living closer to Budongo Forest Reserve reported extensive forest loss outside the reserve since they were children. More recent migrants to the area said the extent of forest cover had not changed since they arrived.

## 4. Discussion

Our conceptual framework brought together three existing frameworks from different literatures in order to understand how interacting with ecosystems may influence psychological distress. In the following, we answer our three research questions by situating our results within the conceptual framework and broader literature. We then describe how these examples help address key gaps in existing evidence linking ecosystems and mental health.

### 4.1. Psychological distress in Nyabyeya

We asked how Nyabyeya’s residents describe their experiences of distress. An extensive literature explores the language people used to express experiences of psychological distress globally (e.g., Kaiser et al., 2015). In addition to indicating psychological distress, these terms can also have symbolic and political connotations, with wide within- and between-cultural variation in meaning (Cork et al., 2019; Kaiser et al., 2015; Pedersen et al., 2010). Consequently, such idioms are unlikely to overlap with psychiatric categories directly. Nevertheless, changes in weight and appetite, sleep disturbance, fatigue and feeling faint, problems concentrating, suicidal ideation, heart palpitations, and chest and abdominal discomfort were associated with ‘thinking too much.’ These symptoms are consistent with a stage-based model of mental illness and can be found along the spectrum of severity (Patel et al., 2018). However, other diagnostic symptoms were not reported by our respondents. For instance, low mood was not mentioned but is indicative of conditions such as depression (APA, 2013). Furthermore, there was variation in the symptoms that respondents associated with ‘thinking too much.’ For instance, some emphasized loss of appetite and weight loss, whereas others highlighted chest discomfort. Consequently, ‘thinking too much’ does not appear to be a single phenomenon. Nevertheless, our results corroborate other studies suggesting that ‘thinking too much’ and related idioms are used to describe psychological distress and poor mental health in Uganda (Kaiser et al., 2015). For instance, a rapid ethnography among perinatal women and healthcare providers in eastern Uganda found overlap between ‘thinking too much’ and diagnostic criteria for major depression, suggesting its association with psychological distress (Tol et al., 2018). When asked about the causes of this distress, many respondents mentioned themes aligned with the Voices of the Poor framework, discussed next.

### 4.2. Social determinants of psychological distress

We explored the perceived stressors contributing to this distress among residents. The following focuses on the most apparent intermediaries between people’s socio-ecological context and experiences of distress (but see Supplementary Information 10: Other pathways). The Voices of the Poor framework suggests that poverty is experienced according to five main themes: *material lack and want*; *physical ill-being*; *bad social relations*; *insecurity and vulnerability*; and *powerlessness, frustration, and anger* (Narayan et al., 2000). The framework stresses that poverty is multidimensional, beyond merely the lack of wealth and income. Yet, respondents’ conception of poverty appeared to be closely associated with a lack of money in our study. Furthermore, the reported experiences of (and responses to) hunger indicate low to moderate food insecurity (Jones et al., 2013). For instance, eating the same food types for multiple meals or eating less preferred foods is a sign of moderate food insecurity. Our results align with those of other studies, which identify poverty and food insecurity as consistent and powerful social determinants of psychological distress and common mental disorders, such as depression (Jones, 2017; Lund et al., 2010; Lund et al., 2018; Weaver & Hadley, 2009). For instance, poverty was a major reported cause of ‘thinking too much’ among residents in central and western Uganda (Miller et al., 2021).

Other themes from the Voices of the Poor framework did not appear to be as salient to the research questions. For instance, physical ill-being appeared to be a source of distress but was not frequently associated with interactions with ecosystems. This result is perhaps surprising, given the many ways that ecosystems are known to affect physical health (Whitmee et al., 2015). For instance, multiple studies associate forest cover and loss with diarrheal disease in tropical countries (e.g., Herrera et al., 2017; Pienkowski et al., 2017). This observation suggests that our approach – which elicits information about links that respondents feel are important and are cognizant of – could be usefully combined with evidence describing less visible connections. For instance, disease ecology research could reveal ways that changing ecosystems might affect locally important physical illness that impacts mental health (e.g., linking deforestation, blackfly populations, and onchocerciasis (Katabarwa et al., 2018)).

In summary, poverty and food insecurity – examples of *material lack and want* and *insecurity and vulnerability* within our conceptual framework – appeared to be the most apparent intermediaries between ecosystems and psychological distress within our case study.

### 4.3. Social determinants within a socio-ecological context

Finally, we asked how interacting with ecosystems, within a wider socio-ecological context, influenced these stressors. Ostrom and colleagues’ framework helped to organize and understand the many features found within Nyabyeya’s socio-ecological context (Ostrom, 2007). Here, we focus on three frequently-described mechanisms (but see Supplementary Information 10: Other pathways).

#### 4.3.1. Agro-ecological systems and psychological distress

Since the early 1990s, Kinyara Sugar Works has expanded its output through the purchase of sugarcane from small-scale contract farmers and sugarcane estates in Nyabyeya (Babweteera et al., 2012; Twongyirwe et al., 2017). Our results indicate that most of those who benefited from this expansion were wealthier families with relatively large amounts of land, many of whom were indigenous Banyoro. Those with less land and capital, many of whom had migrated to the area in the last decades, were less likely earn income through sugarcane cultivation. Moreover, commercial sugarcane farming was reportedly associated with the displacement of subsistence agriculture, exacerbating social determinants of psychological distress among poorer households but alleviating them among wealthier ones (Figure 5). In parallel, migration-driven population growth may also have reduced land availability.

**Figure 5.**
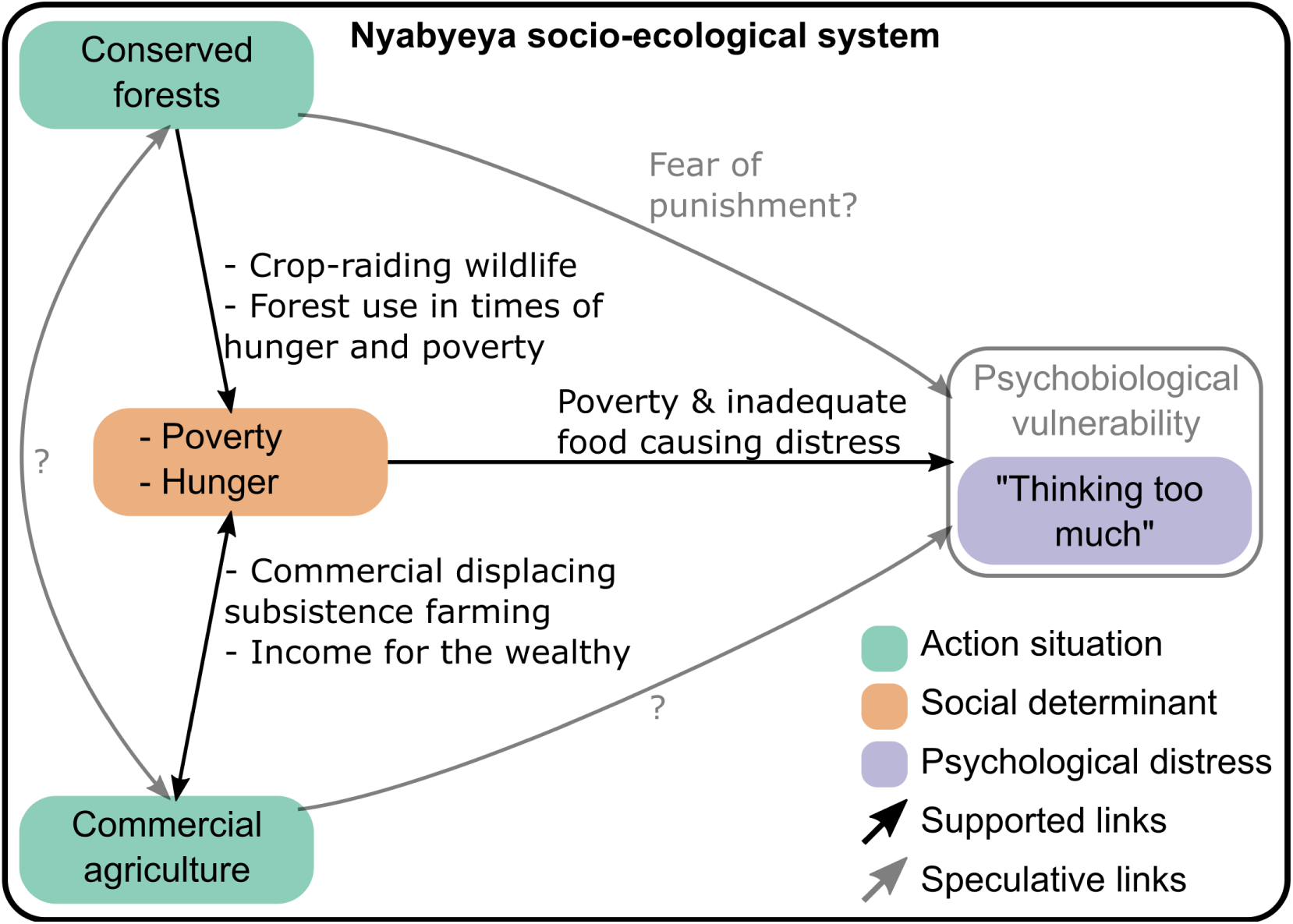
Hypothesized causal connections between two socio-ecological action situations, social determinants, and psychological distress. Interviewee reports substantiate ‘supported’ but not ‘speculative’ links. Commercial agriculture may be a socio-ecological action situation perceived or actually contributing to food insecurity and poverty among subsistence farmers but providing income for wealthier landowners. Simultaneously, those with greater wealth appeared more likely to engage in commercial agriculture. Furthermore, conserved forests may be a socio-ecological action situation supporting crop-raiding wildlife populations while being a source of resources. Crop-raiding is a reported threat to food security. However, illegally and legally harvested forest resources are reported safety nets in the face of poverty or food insecurity. Consequently, conserved forests and conservation management activities may have both desirable and undesirable relationships with poverty and food insecurity. Poverty and food insecurity were reported sources of ‘thinking too much,’ representing possible social determinants of psychological distress.

Recognizing that large-scale agricultural land acquisitions can undermine development efforts, many countries, including Uganda, have promoted contract farming approaches (White et al., 2012). Yet, contract sugarcane farming has also been associated with increased inequality and land dispositional, exacerbating poverty and food insecurity in other parts of Uganda (Martiniello, 2021). For instance, Mwavu et al. (2018) describe how commercial sugarcane farming was associated with reduced availability of land for food crop farming, increasing local food insecurity. More broadly, numerous studies illustrate commercial agriculture’s diverse impacts on smallholder farming systems, food security, and poverty globally (Hall et al., 2017). Therefore, changing food systems could influence social determinants of psychological distress among rural millions globally. Whether these effects are desirable or not may depend on how equitably benefits are distributed, among other factors. This observation demonstrates the value of accounting for wider social systems, such as national policy, when understanding local level mental health.

Industrial agricultural expansion is also a leading cause of environmental degradation worldwide (Lewis et al., 2015). For example, commercial agriculture was one of the largest drivers of forest loss across the tropics between 2001 and 2015 (Curtis et al., 2018). The consequent loss of nature’s contributions could also impact social determinants of psychological distress. A key socio-ecological process that seeks to protect natural systems and the contributions they provide is biodiversity conservation.

#### 4.3.2. Conserved landscapes and psychological distress

Nyabyeya was comparatively sparsely populated when Budongo Forest Reserve was first gazetted in the 1930s (Paterson, 1991). Since then, the population has grown, in part because of in-migration. Simultaneously, there has been a high amount of forest loss outside Budongo Forest Reserve. Much of this loss appears to have occurred before the 2000s, partly attributed to commercial agricultural expansion (Kyongera, 2015; Twongyirwe et al., 2017). In this context, Budongo Forest Reserve appears to be an important contributor to livelihoods, particularly for those experiencing poverty and food insecurity (Figure 5). Our results do not indicate if this importance has changed over time. However, if commercial agricultural expansion has exacerbated food insecurity and poverty among poorer subsistence farmers, then Budongo Forest Reserve may have become an increasingly important safety net for those groups.

Additionally, as no farming is permitted in Budongo Forest Reserve (apart from designed buffer areas), it appears to be an effective barrier to agricultural expansion. This constraint might be an expected source of resentment among those with limited land. However, no respondents expressed this opinion, with some – including those involved in illegal forest use – emphasizing the need to protect the forest from wealthy *“people from outside”* engaged in illegal logging. This result may be because the reserve was gazetted before many families arrived, so might be seen as a fixed feature, while also being a safety net. Yet, not all interactions with conserved forest ecosystems were desirable, as discussed next.

Crop-raiding was associated by respondents with crop losses and the disruption of income-generating activities, both reportedly contributing to food insecurity and poverty, and consequent distress. However, the forest-farm frontier has reportedly shifted over time, so that it now coincides with the boundaries of the two forest reserves. Crop-raiding by wildlife is likely to have always affected households at the forest edge. However, it appears that both the location of the forest edge and the groups living at it have changed over time, with migrants tending to settle in available land at the forest-farm frontier and indigenous Banyoro typically found closer to community centers. This emphasizes how the psychological impacts of interactions with ecosystems may affect different groups over time, as the result of changing social and environmental contexts.

In general, nature conservation seeks to maintain ecosystems, with protected areas covering 15% of low- and middle-income countries’ terrestrial areas (UNEP-WCMC et al., 2018). The protection of nature’s contributions may, therefore, influence social determinants of psychological distress among millions worldwide. For instance, small-scale wild-capture fisheries support the livelihoods of over 100 million people in developing countries (World Bank et al., 2012). Sustainable management of fisheries may help sustain these livelihoods and, indirectly, mental health. Yet, not all interactions with ecosystems are desirable. Several studies describe how interactions with wildlife can be distressing, potentially contributing to the risk of mental illness (e.g., Barua et al., 2013; Chowdhurym et al., 2016; Jadhav & Barua, 2012). For instance, Chowdhurym et al. (2016) interviewed 65 widows whose husbands had been killed by tigers, and who subsequently faced stigma and other social challenges. Their study suggested that respondents’ status as *“tigers-widows”* contributed to their risk of mental illness, with 44% of them being formally diagnosed with a mental disorder.

Furthermore, conservation areas themselves can have varied material and non-material impacts on people’s well-being (Thondhlana et al., 2020; Woodhouse et al., 2015). For example, there are multiple instances where residents have been displaced and impoverished through the management of conservation areas (Agrawal & Redford, 2009). Equally, there are instances where protected areas and associated management strategies contribute to residents’ well-being (e.g., Naidoo et al., 2019). Conservation scientists recognize that well-being is multidimensional and includes both health and subjective evaluations of well-being (Woodhouse et al., 2015). However, little research appears to explore the roles of conservation in psychological distress and mental illness. For instance, McKinnon et al. (2016) systematically reviewed 1043 articles and grey literature exploring the impacts of conservation on human well-being, identifying 60 that focused on health. We examined the abstracts and summaries of these and found just one that focused on the relationships between conservation and psychological distress or mental illness. This one exception was a study describing the psychological trauma associated with human-wildlife conflict around conservation areas in India (Ogra, 2008).

Despite the limited number of articles explicitly connecting conserved ecosystems and mental health, we believe these links are prevalent and important. For instance, McKinnon et al. (2016) also found 700, 310, and 94 articles relating conservation to economic living standards, social relations, and security and safety, respectively, all of which are recognized social determinants of common mental disorders (Lund et al., 2018). Given this, it is highly likely that conservation has myriad desirable and undesirable impacts on people’s lives in ways that indirectly influence the risk of psychological distress and potentially mental illness. Given the magnitude of the global burden of mental illness and the extent to which conservation influences lives worldwide, this appears to be a significant omission from conservation research.

### 4.4. Policy relevance of addressing gaps in the evidence base

We identified two gaps in understanding how interacting with ecosystems influences psychological distress and potentially mental illness; the neglect of the rural global south and overlooking the intermediate role of social determinants.

First, we used the shorthand ‘global south’ to mean regions of Africa, Asia, Oceania, and Latin America. However, the use of this term risks clustering the diverse ways that interacting with ecosystems may influence psychological distress across countries and contexts. For instance, within our case study, the psychological impacts of changing agricultural systems appeared to be different between wealthier and poorer households. Likewise, this clustering obscures the many similarities in experiences between those in the global north and south. For instance, the psychological costs of failed harvests might be similar for farmers in rural Uganda and England. Consequently, while our case study illustrates new mechanisms from a global south perspective, it also suggests the need for a locally nuanced understanding of links in different socio-ecological contexts more generally. The diversity of these links should be reflected in policymaking at the interface between ecosystems and health. For instance, the *Zero Draft of the Post-2020 Global Biodiversity Framework* suggests enhancing access to green spaces for health and well-being (CBD, 2020). This target might be desirable in some contexts, but our study points to other ways that environmental policymakers can indirectly support mental health. For instance, promoting equitable resource access and control through participatory landscape governance may create conditions supporting mental health. Generally, many of the ways that ecosystem management might influence mental health are likely to be indirect, as discussed next.

Second, *The* Lancet *Commission on global mental health and sustainable development* situated mental well-being within the global Sustainable Development Goals (SDGs, Patel et al., 2018). The *Commission’s* report drew on a comprehensive review of the literature describing the social determinants of mental illness, which included the role of natural disasters and climate change (Lund et al., 2018). However, links with ecological SDGs such as those related to life on land and in water were largely absent, perhaps because of the limited research linking these topics. Given the lack of progress towards ecological goals known to influence human well-being and the recognized importance of nature’s contributions, this omission may miss valuable policy opportunities (IPBES, 2020; Shepherd et al., 2016). Specifically, there may be opportunities to foster co-benefits between SDG 3, related to good health and well-being, and other SDGs. For instance, sustainable fisheries that support life in water (SDG 14) can help end hunger (SDG 2), thereby perhaps also alleviating stressors contributing to mental illness (SDG 3, Nash et al., 2020). Equally, there are also likely to be detrimental trade-offs between mental health and other aspects of the SDGs, which ought to be managed. For instance, although modernizing agricultural ecosystems can help meet many SDGs, it can also compound inequality (SDG 10) and hunger (SDG 2) for some, potentially exacerbating risks to mental health (SDG 3, Dawson et al., 2019).

Understanding these indirect co-benefits and trade-offs may also help make the case for protecting some types of ecosystems. The ‘New Deal for Nature’ seeks to mainstream the valuation of biodiversity into political, economic, and social systems (UNEP, 2019b). Some of the direct ways that ecosystems can support mental health have been valued in monetary units easily integrated into policymaking. For instance, Buckley et al. (2019) estimate the global value of improved mental health as the result of visiting green spaces in protected areas at US$6 trillion per year. Yet, to our knowledge, the monetary values associated with indirect pathways between ecosystems and mental health have not been estimated but are likely to be substantial. Recognizing and accounting for the value of these indirect pathways would add impetus for protecting ecosystems within the ‘New Deal for Nature.’

Furthermore, investigating the roles of socio-ecological context could also help explain the etiology of mental illness. On the one hand, this may offer new opportunities to help tackle the *“global tragedy”* of avoidable mental illness (Becker & Kleinman, 2013, p. 66). On the other hand, it may also reveal how unchecked ecological degradation may worsen the global burden of mental illness. In both cases, this better understanding may also help foster cooperation between public health and environmental experts seeking to address shared goals.

## 5. Conclusion

The global mental health and ecological crises present growing threats to human well-being (IPBES, 2020; Patel et al., 2018). While an increasing number of studies explore how changes to ecosystems harm physical health, relatively few explore ecosystems’ roles in mental illness. We use a new framework that situates social determinants of psychological distress in socio-ecological systems. Using this framework, we show that commercial agriculture and nature conservation – land uses affecting the lives of millions – may affect social determinants of psychological distress in our case study. Further research is required to understand if these links with psychological distress are also associated with mental illness.

Nevertheless, accounting for people’s interactions with ecosystems may help better explain the etiology of mental illness, particularly if ecological change continues to escalate. Furthermore, our study reveals ways to potentially manage socio-ecological systems to support mental health, such as promoting fair access and control of natural resources. Finally, our study indicates opportunities to manage co-benefits and trade-offs between sustainable development goals, which may also support mental health. In summary, our case study demonstrates the importance of taking a more holistic and locally nuanced approach to understanding how people’s relationships with ecosystems relate to psychological distress and mental illness.

## 6. Acknowledgments

TP would like to thank the participating residents of Nyabyeya, Moses Musiimenta, Susan Lajiki, and Liberty Anichan for their assistance with the study, and Ruth Pinto for her comments. This work was supported by the Natural Environment Research Council (grant number NE/L002612/1), the Parkes Foundation Small Grant Fund, Wolfson College at the University of Oxford Travel Grant, and the Africa-Oxford Initiative Travel Grant. The funders had no involvement in study design; in the collection, analysis, and interpretation of data; in the writing of the report; and in the decision to submit the article for publication.

## 7. Author contributions

Thomas Pienkowski: conceptualization; data curation; formal analysis; funding acquisition; investigation; methodology; project administration; resources; software; validation; visualization; roles/writing - original draft; writing - review & editing.

Aidan Keane: conceptualization; formal analysis; funding acquisition; methodology; resources; supervision; validation; roles/writing - original draft; writing - review & editing.

Eugene Kinyanda: conceptualization; formal analysis; funding acquisition; methodology; resources; supervision; validation; roles/writing - original draft; writing - review & editing.

Caroline Asiimwe: conceptualization; formal analysis; funding acquisition; methodology; resources; supervision; validation; writing - review & editing.

Geoffrey Muhanguzi: conceptualization; formal analysis; funding acquisition; methodology; resources; supervision; validation; writing - review & editing.

Birthe Loa Knizek: formal analysis; funding acquisition; methodology; resources; supervision; validation; writing - review & editing.

E.J. Milner-Gulland: conceptualization; formal analysis; funding acquisition; investigation; methodology; project administration; resources; supervision; software; validation; visualization; roles/writing - original draft; writing - review & editing.

