## Supplementary Information for "The role of nature conservation and commercial farming in psychological distress among rural Ugandans"

### S1: Article search method

Two searches were performed on the 14th June 2020 in the Scopus abstract and citation database (Scopus, 2020). The first uses the following search string:

> (TITLE-ABS ("natural environment")) AND (TITLE-ABS (("human" OR "people" OR "humanity"))) AND (TITLE-ABS (("health") OR ("illness") OR ("disease"))) AND (LIMIT-TO (DOCTYPE, "ar"))

This first search returned 1,306 results. The second included additional search terms associated with common mental disorders (NCCMH 2011):

> (TITLE-ABS ("natural environment")) AND (TITLE-ABS (("human" OR "people" OR "humanity"))) AND (TITLE-ABS (("health") OR ("illness") OR ("disease"))) AND (TITLE-ABS (("mental health") OR ("mental illness") OR ("mental disorder") OR ("depression") OR ("depressive disorder") OR ("anxiety") OR ("post-traumatic stress") OR ("panic disorder") OR ("social anxiety") OR ("obsessive-compulsive disorder") OR ("phobias"))) AND (LIMIT-TO (DOCTYPE, "ar"))

This second search returned 65 results or 5% of the first search.

### S2: Conceptual framework

The conceptual framework has two purposes. First, to illustrate plausible ways by which interacting with ecosystems could influence psychological distress. Second, to guide the semi-structured interviews within the case study.

Multiple frameworks illustrate the ways that natural processes directly affect health (e.g., Bayles et al., 2016; Myers et al., 2013; Myers & Patz, 2009; Sandifer et al., 2015). However, since social determinants may mediate many links between ecosystems and psychological distress, we adopt a framework that explicitly illustrates the role of social determinants (Lawrence et al., 2019). Berry et al. (2018) provide such a framework in the contact of climate change. However, our case study is interested in a broad range of socio-ecological dynamics, and so we adopt a framework with three components (Figure 1).

The framework argues that socio-ecological systems define the context in which people live their lives (Fisher et al., 2013), understood using a socio-ecological systems framework (McGinnis & Ostrom, 2014; Ostrom, 2007, 2009). These stressors can be structured into major domains using the 'Voices of the Poor' framework (Narayan et al., 2000). Excessive exposure to stressors, in the form of chronic strains, "daily hassles," and major life events may increase the risk of poor mental health (Serap  Keles et al., 2016; Lamis & Kaslow, 2014; McIntosh et al., 2010; Ormel & Neeleman, 2000; P. Spinhoven et al., 2010; Thoits, 2010; Zubin & Spring, 1977). The following provides further details on each step within this framework.

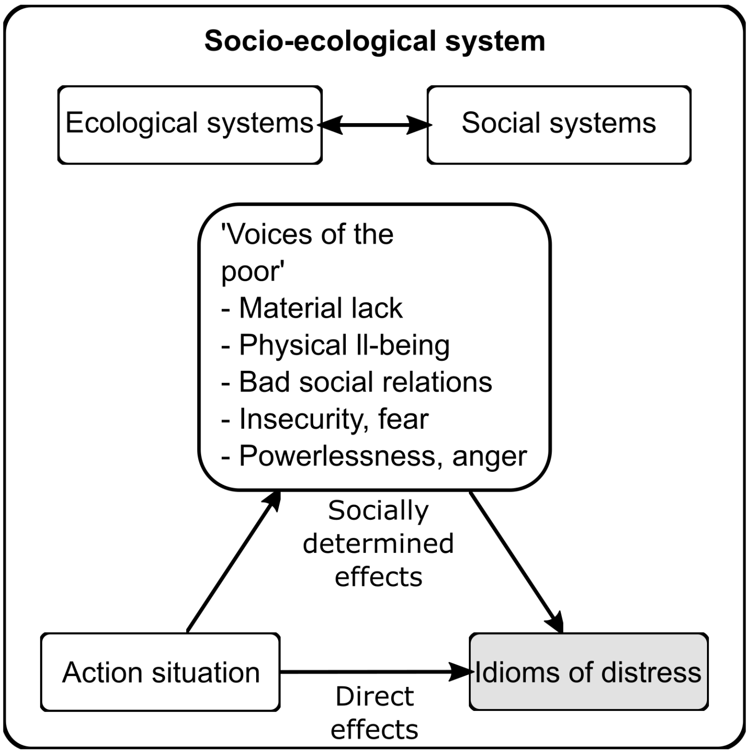


Figure 1. Illustrating how 'action situations' within socio-ecological systems might influence stressors faced by populations experiencing poverty, using the Voices of the Poor framework. These stressors can be considered as social determinants of psychological distress, and accompany direct relationships between socio-ecological systems and psychological distress. This psychological distress can be expressed in locally relevant idioms.

### The psychological distress framework

The first component describes how the experience of stressors over an individuals' life-course can contribute to psychological distress, depending on an individual's psychobiological vulnerability (Goh & Agius, 2010; Johnstone et al., 2018; Patel et al., 2018). Stress has been described as a process by which *"environmental demands tax or exceed the adaptive capacity of an organism, resulting in psychological and biological changes that may place persons at risk for disease"* (Cohen et al., 1995). Although some degree of stress can be beneficial, distress occurs at levels of stress that are harmful to mental health (Le Fevre et al., 2003; Ridner, 2004).

Stressors can vary in their intensity and frequency; stressors are sometimes considered in terms of major life events, *"daily hassles,"* and chronic strains (Kanner et al. 1981; Pearlin 1989). There is now significant evidence that the experience of negative life events, particularly during childhood and adolescence, increases risks to mental health (Penninx et al. 2010). Furthermore, the effect of stressors may be cumulative and persist over long periods (Seery 2011; Seery, Holman, and Silver 2010). Studies have shown that stressors in the form of daily hassles and chronic strains may be important risk factors for poor mental health (Serap Keles et al., 2016; Lamis & Kaslow, 2014; McIntosh et al., 2010). Importantly, an individual's risk of poor mental health, and the pathogenic effect of stressors, also depends on their genetic, neurodevelopmental, and psychological characteristics (Kessler & Bromet, 2013; Lund et al., 2018; Philip Spinhoven et al., 2010).

More generally, an individual's movement between mental health and illness is increasingly seen as an evolutionary process in which distress is a prevalent feature (McGorry et al., 2014; Patel et al., 2018). This stage-based model of mental illness suggests a spectrum from mental well-being, through diffuse and non-specific psychological distress, to definable syndromes (McGorry et al., 2014; McGorry & van Os, 2013). At the early stages, individuals may experience a wide range of non-specific symptoms, such as fatigue, anxiety, or insomnia. At later stages, these can become more severe, specific, and resistant to treatment, eventually passing diagnostic thresholds for mental illness (Patel et al., 2018). Individuals can move in both directions along this spectrum, and the experience of stressors over an individuals' life-course can be risk factors within this stage-based model (Johnstone et al., 2018; Lund et al., 2018). Here, we recognize that individuals can move between stages of mental health and illness, and psychological distress may be experienced throughout this process (Drapeau et al., 2012; Johnstone et al., 2018; McGorry et al., 2014; Patel et al., 2018).

Classification systems such as the Diagnostic and Statistical Manual of Mental Disorders simplify experiences of poor mental health into discrete diagnosable disorders (APA 2013). Although offering advantages, such diagnostics can be overly reductionist, particularly when exploring cross-cultural experiences of mental illness (Johnstone et al., 2018). We explore experiences of distress using locally appropriate terms; these idioms of distress are *"socially and culturally resonant means of experiencing and expressing distress in local worlds"* (Nichter, 2010). Idioms of distress may be used by individuals experiencing varying degrees of mental health and illness, can have symbolic and political meaning, and are culturally situated (Cork et al., 2019; Kaiser et al., 2015; Pedersen et al., 2010).

### The Voices of the poor framework

The second component describes broad categories of stressors faced by populations experiencing poverty, some of which may represent social determinants of mental illness. This component draws on the 'Voices of the Poor' initiative, a global effort by the World Bank to understand the experiences, priorities, aspirations, and reflections expressed by poor people themselves (Narayan et al., 2000). The project used participatory methods, with more than 20,000 individuals in 24 countries. The emergent findings can be considered a framework for understanding the multiple dimensions of ill-being and are used in this project to identify broad types of stressors within residents' lives.

The framework makes the distinction between well-being and ill-being, which included five main domains. Firstly 'material lack and want,' which included food, livelihood, assets and money, and housing and shelter. Secondly, 'physical ill-being,' which included hunger, pain and discomfort, and exhaustion and poverty of time. Thirdly, 'bad social relations: exclusion, rejection, isolation, and loneliness.' Fourth, 'insecurity, vulnerability, worry, and fear.' Finally, 'powerlessness, helplessness, frustration, and anger.' Here we focus on ill-being domains since these are expected to represent stressors that may emerge within a socio-ecological context.

### The socio-ecological systems framework

The final component describes how socio-ecological systems define the context of people's lives, including the stressors that they experience. Socio-ecological systems are *"interdependent and linked systems of people and nature that are nested across scales"* (Bouamrane et al., 2016). Ostrom and colleagues provided a framework for organizing and structuring the many variables found in socio-ecological systems within a nested framework (Colding & Barthel, 2019; McGinnis & Ostrom, 2014; Ostrom, 2007, 2009). This socio-ecological systems framework is typically used to understand natural resource use (McGinnis & Ostrom, 2014; Partelow, 2018). However, the framework can be used more broadly, such as helping conservation planners understand and manage social processes that influence conservation outcomes (Ban et al., 2013).

The socio-ecological systems framework includes subsystems resource units (e.g., trees, crops), resource systems (e.g., a conserved landscape), governance system (e.g., a local council, a conservation NGO), and actors (e.g., farmers, children, conservation staff, Figure 2). The interactions of these subsystems generate outcomes: those interactions and outcomes of most interest to a user are termed 'action situations.' The broader social (including political and economic) setting and related ecosystems dynamically interact with elements of the core subsystems. We use the socio-ecological system's framework as a tool to think systematically about the social and environmental domains that may be associated with specific stressors.

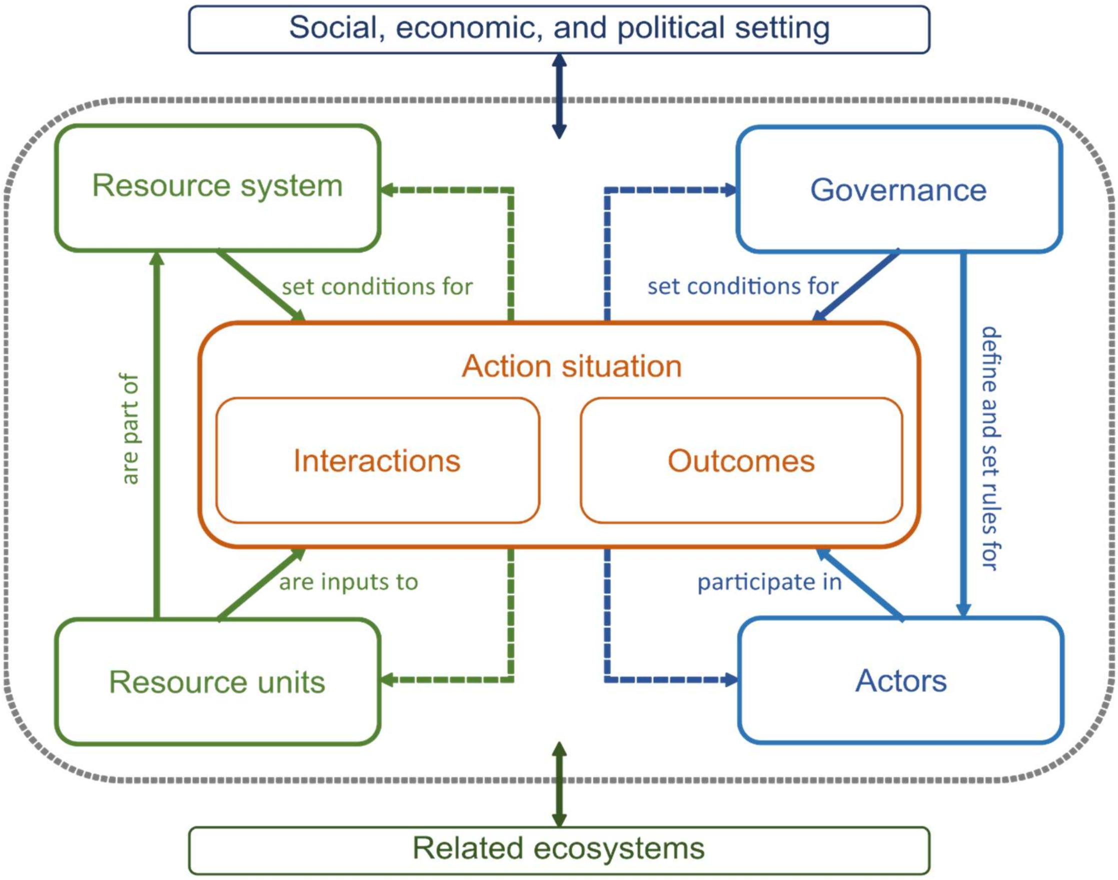


Figure 2. The socio-ecological system framework, adapted from (Ostrom, 2007).

### S3: Ethical considerations

The ethical procedure was developed after reviewing the *Code of Conduct for researchers contributing articles to Oryx – The International Journal of Conservation* (Oryx, 2001). Ethical approval was granted by the Uganda National Council of Science and Technology (Ref. SS6007), Government of Uganda, and the the Central University Research Ethics Committee (Ref. R63458) at the University of Oxford.

Before commencing the interviews, respondents were informed of the purpose of the research and who was conducting it. They were also informed that participation was voluntary, and respondents could withdraw anytime during or after the interview. Respondents were informed about how the data would be used, the risks of participation, the complaints procedure, and contact details for the ethical review board that approved the study. Documented, free, prior, and informed consent was then sought before individuals could take part. Any respondents that reported that they were under the age of 18 were not interviewed.

Respondents were debriefed after completing the interview, which included providing the contact details of the research team, and the Ethical Review Board. Respondents were provided with household essentials (including soap, salt, oil, and sugar) as in-kind compensation for their time.

### S4: Sampling effort

Table 1. The sampling effort in each community, and approximated gender, age, socioeconomic status, and forest proximity. Socioeconomic status was subjectively judged, based on indicators suggested to us by the Local Council members, including the type of house construction, cooking and toilet facilities, presence of assets such as cars, chairs, etc. TC = Trading Centre.

| Community | Gender | Age | Socioeconomic status | Forest proximity | Study ID |
|---|---|---|---|---|---|
| Nyabyeya TC | Female | Middle-aged | Middle | Far | 1 |
| Nyabyeya TC | Male | Middle-aged | Middle | Far | 2 |
| Nyabyeya TC | Male | Middle-aged | Low | Closer | 3 |
| Nyabyeya TC | Female | Middle-aged | High | Far | 4 |
| Nyabyeya TC | Female | Older | Low | Closer | 5 |
| Nyabigoma | Male | Older | High | Far | 6 |
| Nyabigoma | Female | Older | Low | Far | 7 |
| Nyabigoma | Female | Young | Middle | Closer | 8 |
| Nyabigoma | Male | Middle-aged | Middle | Far | 9 |
| Nyabigoma | Female | Young | Middle | Far | 10 |
| Nyabigoma | Male | Young | Middle | Far | 11 |
| Nyakafunjo | Male | Young | Middle | Adjacent | 12 |
| Nyakafunjo | Male | Young | Middle | Adjacent | 13 |
| Nyakafunjo | Male | Older | Middle | Adjacent | 14 |
| Nyakafunjo | Female | Young | Low | Adjacent | 15 |
| Nyakafunjo | Female | Young | Low | Adjacent | 16 |
| Kyempunu | Male | Young | Middle | Adjacent | 17 |
| Kyempunu | Female | Middle-aged | Middle | Adjacent | 18 |
| Kyempunu | Male | Young | High | Far | 19 |
| Kyempunu | Female | Older | Middle | Closer | 20 |
| Kyempunu | Male | Middle-aged | Middle | Adjacent | 21 |
| Nyabyeya 2 | Male | Young | Middle | Closer | 22 |
| Nyabyeya 2 | Female | Older | Low | Closer | 23 |
| Nyabyeya 2 | Female | Middle-aged | High | Adjacent | 24 |
| Nyabyeya 2 | Male | Middle-aged | Middle | Far | 25 |
| Nyabyeya 2 | Female | Middle-aged | Middle | Closer | 26 |
| Kanyege | Female | Middle-aged | Low | Far | 27 |
| Kanyege | Female | Middle-aged | High | Far | 28 |
| Kanyege | Male | Middle-aged | Low | Far | 29 |
| Kanyege | Female | Young | Low | Closer | 30 |
| Karongo | Female | Middle-aged | High | Closer | 31 |
| Karongo | Male | Middle-aged | High | Adjacent | 32 |
| Karongo | Female | Middle-aged | Low | Closer | 33 |
| Karongo | Male | Middle-aged | Middle | Closer | 34 |
| Karongo | Male | Middle-aged | Low | Far | 35 |
| Maramu | Female | Middle-aged | High | Far | 36 |
| Maramu | Female | Middle-aged | Low | Adjacent | 37 |
| Maramu | Male | Older | Middle | Closer | 38 |
| Maramu | Male | Middle-aged | High | Closer | 39 |
| Maramu | Male | Middle-aged | Middle | Adjacent | 40 |
| Nyabyeya 1 | Female | Older | Middle | Far | 41 |
| Nyabyeya 1 | Female | Older | Middle | Far | 42 |
| Nyabyeya 1 | Female | Middle-aged | High | Far | 43 |
| Nyabyeya 1 | Female | Young | Middle | Closer | 44 |
| Nyabyeya 1 | Male | Older | Middle | Closer | 45 |

### S5: Interview guide

#### [*Actors*]: Tell me about the people that live in this area?

- History of the community and people within it
- Leaders
- Community groups, other groups
- Socioeconomic attributes
- Gender
- Ethnicity
- Age

[***Resource systems***]: Tell me about the important places in and around the community?

- Important places
- Important resources
- Access to what areas/resources
- Human facilities
- Productivity of system
- Seasonality and variation over time

[***Resource units***]: What are the most important resources in your daily lives?

- Resources
- Location of resources
- Abundance or size of the resource
- Wildlife
- Plants
- State of resources
- Change over time

[***Interactions and Outcomes***]: What do people do in the area?

- Livelihoods
- Food
- Fuel
- Water
- Medicine
- Spiritual and cultural
- Recreation
- Harvesting
- Information sharing
- Conflicts
- Deliberation and decision
- Investment
- Lobbying
- Networks and social relations

[***Governance systems***]: How are decisions made about the community and environment?

- Laws and rules
- Governance organizations
- Access and property rights
- Informal and formal decision making
- Civil society organizations
- Relations between actors and systems
- Sanctions

[***Social, economic, and political settings***]: How does the wider economy influence the community?

- Economic development
- Business
- National government

[***Related ecosystems***]: Changes in the wider world?

    - Climate change

#### ‘VoP’

##### *Material lack and want*

[***Material lack and want***]: What problems do people have about having enough in life?*

- Food
- Livelihood, assets, and money
- Housing and shelter

[***Vulnerability***]: Who is most affected by these problems?‡

[***SESF-link***]: Why do people have these problems?

##### *Physical ill-being*

[***Physical illbeing***]: What problems do people have with their health?*

- Hunger, pain, and discomfort
- Exhaustion and poverty of time

[***Vulnerability***]: Who is most affected by these problems?‡

[***SESF-link***]: Why do people have these problems?

#### *Bad social relations*

[***Bad social relations***]: What problems do people have with their relationships with others?*

[***Vulnerability***]: Who is most affected by these problems?‡

[***SESF-link***]: Why do people have these problems?

#### *Insecurity, vulnerability, worry, and fear*

[***Insecurity, vulnerability, worry, and fear***]: What are people worried about and what causes them fear?*

[***Vulnerability***]: Who is most affected by these problems?‡

[***SESF-link***]: Why do people have these problems?

#### *Powerlessness, helplessness, frustration, and anger*

[***Powerlessness, helplessness, frustration, and anger***]: What makes people feel angry or like they have no control in their lives?*

[***Vulnerability***]: Who is most affected by these problems?‡

[***SESF-link***]: Why do people have these problems?

### Secondary questions

*To be asked in relation to major stressors identified in the interview*

*[***Idioms of distress***]: What feelings do people have when they have these problems?

*[***Stressor characteristics***]: How long do these feelings last, and how often do they happen?

‡[***Factors affecting stressor exposure***]: Why are these people most affected by these problems?

### S6: Steps in the thematic analysis

We employed inductive thematic analysis identifying, analyzing, organizing, and reporting patterns of themes within data through the following steps, following Braun and Clarke (2006).

1. Familiarization with data: Two research assistants who lived in the study communities and TP discussed the themes and topics immediately after each interview, which were documented in a post-script. TP read each transcript, as well as compared word clouds between different groups (based on gender, apparent socioeconomic status, age, and proximity to the forest edge).
2. Generating codes and coding text: Two sets of codes were developed by TP. The first was created following the initial round of familiarization. These five codes were broad and used to cluster related text. The second set of codes was developed by reading the clustered text. These were then systematically applied by TP to all transcripts through a second round of coding the clustered text. In some cases, codes were revised (split or combined), and new codes identified during the second round of coding.
3. Searching and clustering into themes: Themes were identified by TP by clustering related codes. Some themes emerged during the development of the second set of codes. Other themes were identified when reviewing text within each code. These themes were explored by visualizing connections between codes as a network. Specifically, for each interview, any given section of the text may have been given multiple codes. When a section of text was coded as two more codes, then this represented a connection between those codes. A maximum of one connection was recorded for each interview for any given pair of codes. These connections are represented as a network for selected themes of interest. Within the network, the width of the connector correspondents to the number of interviews reporting the connection. The size of the nodes corresponds to the number of interviews mentioning that code. For instance, if a section of text was coded under codes' A' and 'B,' then a connection between the two was recorded for that interview. A maximum of one connection was recorded for any two codes in an interview. These were then visualized as network diagrams in the 'R' programming language, using the package 'igraph' (Csárdi & Nepusz, 2006; R Core Team, 2020).
4. Reviewing themes: The codes and related text was reviewed to ensure that there was consistency within the theme, but discrete differences between themes. In some cases, themes were merged or split apart.
5. Defining and naming themes: A short description was provided for each theme, by TP.

### S7: Definition of key themes

| Theme name | Definition |
|---|---|
| [Symptoms of] "Thinking too much" | The physical and psychological 'symptoms' experienced when "thinking too much," illustrated in the main text (Figure 3). |
| [Causes of] "Thinking too much" | The causes of "thinking too much," illustrated in the main text (Figure 4). |
| Causes of poverty | The causes of poverty. Poverty often used synonymously with not having enough money but also as a more general state defined by multiple deficiencies. |
| Farm productivity | The factors influencing subsistence and small-scale commercial farming. |
| Causes of not enough land | The explanations for why respondents said there was not enough land. |
| Sugarcane displacing crop farming | The reasons why respondents said that sugarcane farming was perceived to be displacing sugarcane farming (although many said this was happening, without providing a reason). |
| Drivers of development | The factors that bring development to a household. |
| Consequences of poverty | The consequences of experiencing poverty. |
| Causes of hunger | The causes of hunger, or not having enough food. |
| Crops losses or poor harvests | The causes of crop-losses and poor harvests. |
| Crop-raiding consequences | The consequences of crop-raiding by wildlife, for affected households. |
| Consequences of hunger | The consequences and coping strategies associated with hunger or not having enough food. |
| Using the forest | The ways that people used the forest. |
| Illegal forest use risks | The risks associated with engaging in illegal forest activities. |

### S8: Positionality statement

The following describes my (TP) positionality as the lead author within the research. I am a straight white young and educated male who grew up in a middle-class politically left-leaning family in the United Kindgom. I am not religious or spiritual, have socially and economically progressive views, and have worked and studied in nature conservation for ten years. Within the case study, I would be positioned as an outsider, with many of my characteristics contrasting sharply with residents. These are likely to influence multiple aspects of the project, as described below.

The project was conceptualized and developed primarily by myself, EJMG, AK, and EK. As a result, the choice of research questions, the conceptualization of the problem, and the way the research was approached were mostly developed by those least similar to Budongo's residents. This research is largely extractive, with most of the direct benefits accruing to the academics, but with the intention that the results are useful to local conservation (such as Budongo Conservation Field Station) and residents.

I led the semi-structured interviews, with the assistance of two women who came from the study communities, who translated the dialogue. A Ugandan male from outside the study communities also assisted with logistics and interviews. There are several key implications of my positionality in the interviews:

- Some residents expected that, because an outsider with my characteristics was leading the research, it would lead to a project that might benefit them, despite the pre-interview information telling them otherwise. As a result, some respondents may have exaggerated the challenges they face in the expectation of future gain.
- Residents within one community were concerned that I might have been there to appropriate their land. This concern was because they had an ongoing land dispute with a local estate owner, whom residents suspected I might have been working for. Consequently, residents in this community appeared hesitant to discuss land issues openly.
- Relatedly, residents probably withheld sensitive information, including regarding illegal behaviors (such as logging in the forest) or witchcraft. Whereas illegal behaviors were not central to the research, witchcraft plays an important role in the causes and experiences of illness. When discussing witchcraft, residents would sometimes report the experiences or beliefs of others, rather than themselves. Furthermore, being non-religious and non-spiritual may have limited my understanding and, therefore, my ability to investigate these topics.
- Being a male also likely influenced what I was told, and my ability to investigate gendered topics. For example, women were unlikely to have disclosed experience of accessing

contraception or items used during periods. Although potentially prevalent, issues of sexual violence were only raised by a few respondents.

- HIV/AIDS is stigmatized, and residents reportedly did not openly discuss it. The three people I worked with during the interviews pointed out that during interviews, residents would intimate that they had HIV/AIDS, and how it affected them, without openly disclosing it.
- I was told that people would only talk about "too many thoughts" among close friends and family. However, the topic of "too many thoughts" was elicited unprompted by a relatively large number of respondents, when asked how a given stressor affected them. This may be because discussing the topic with a stranger had relatively low social costs, with the belief among some that I would help them.
- An educated white male has high social status, with consequent power imbalances. As a result, this may have increased the chance that respondents were guided by accidental leading questions or being unwilling to lead the conversation.

### S9: Other pathways

The main text presented themes relating interactions with ecosystems to experiences of distress that were most consistently mentioned by respondents. However, there was some evidence of other potential pathways for how interacting with ecosystems may influence social determinants of psychological distress.

1. One mechanism by which land reportedly transitioned from subsistence to small-scale commercial farmland was through the voluntary sale of land. When asked why respondents sold land, several said this was a means of getting money to meet immediate needs. These immediate needs included paying for healthcare, school fees, or for house construction. However, a few suggested men would sell land to pay for alcohol, gambling, or during affairs. All of these were mentioned as contributors to household conflict, as well as food insecurity, and therefore potentially psychological distress.
2. Several respondents said that water sources within the two forest reserves were better quality than those outside the reserves, in farmland. A few respondents said that water quality affected the risk of diarrheal disease and other illnesses, affecting physical health. Physical health was a major stressor associated with psychological distress. However, the physical illnesses associated with distress were typically major disabilities, chronic illness, and HIV/AIDs – suffering diarrheal disease itself did not appear to be a major source of distress.
3. Many female respondents reported fear of chimpanzees, particularly when they were working on farms close to the forest edge or when collecting firewood from Budongo Forest Reserve. This fear was not a reported cause of psychological distress. However, during data collection, a young child who had been taken into the forest by their mother was severely injured by a chimpanzee. Such events are rare, but one of the research assistants said the family subsequently left the community because of stigmatization. This may have represented a source of distress for this family.
4. Several respondents said that two local NGOs and the National Forestry Authority were intending to reforest buffer strips within river valleys. Several of these respondents claimed that riparian buffer areas were under the jurisdiction of the National Forestry Authority. However, some households were farming on these areas and were informed they would be removed from riparian buffer land, which was to be subsequently reforested. This did not appear to have commenced at the time of data collection. However, given concerns about land availability and its possible role in poverty and food insecurity, this may be a source of distress.

### S10: Perceived causes and consequences of poverty

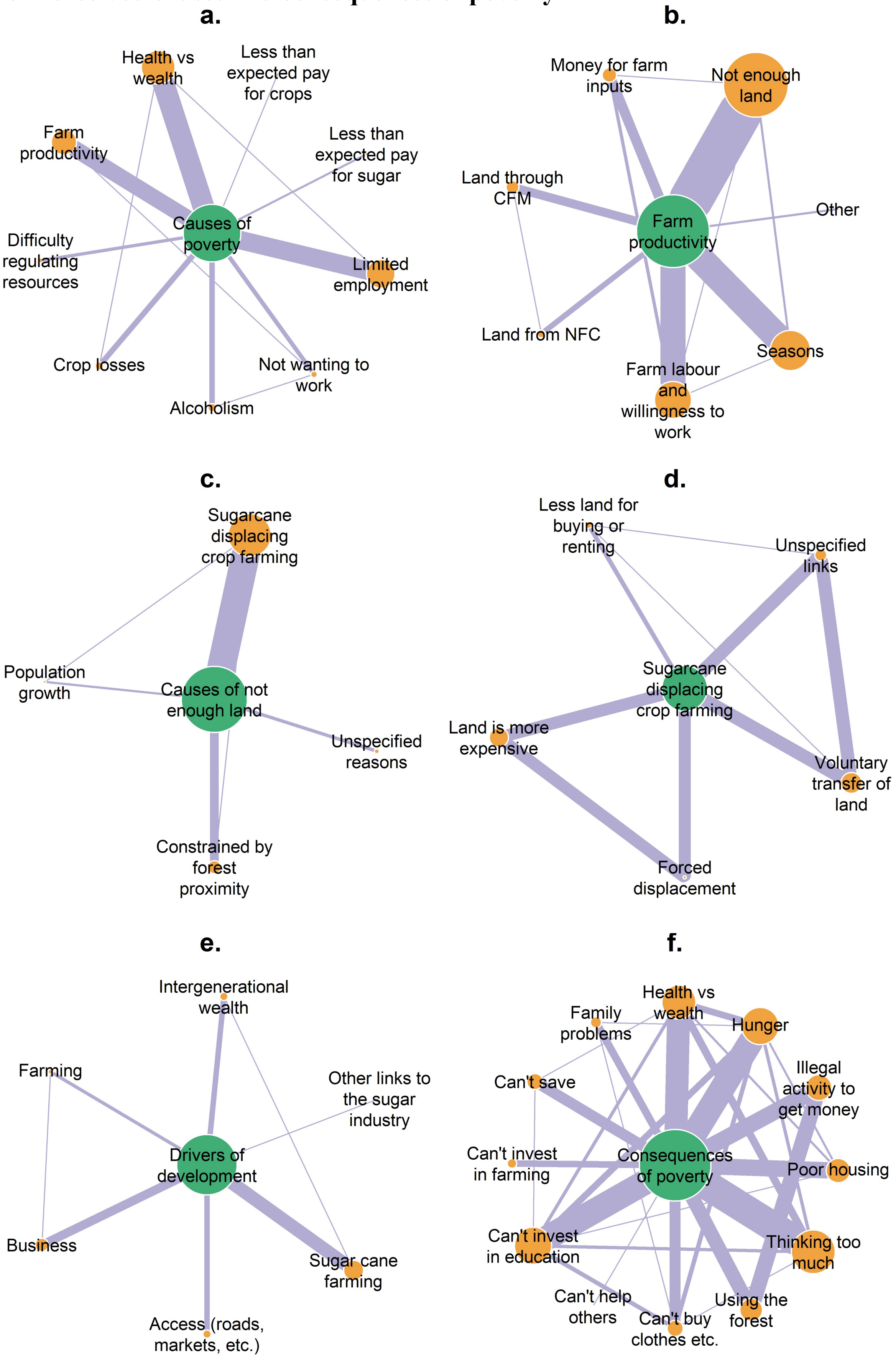

Figure 3. The width of the lines illustrates the number of interviews that reported connections between nodes but did not imply the direction of the causality of connections. The size of the node represents the number of interviews mentioning the associated theme for that node. Panel a. illustrates the reported causes of poverty and inadequate money. Panel b. illustrates reported factors affecting farm productivity (CFM = Community Forest Management, NFC = Nyabyeya Forestry College). Panel c. describes the reported causes of inadequate land. Panel d. illustrates the reported role of the sugarcane industry in displacing smallholder farming. Panel e. describes reported factors that contributed to household development. Panel f. illustrates the reported consequences of poverty and inadequate money.

### S11: Perceived causes and consequences of hunger

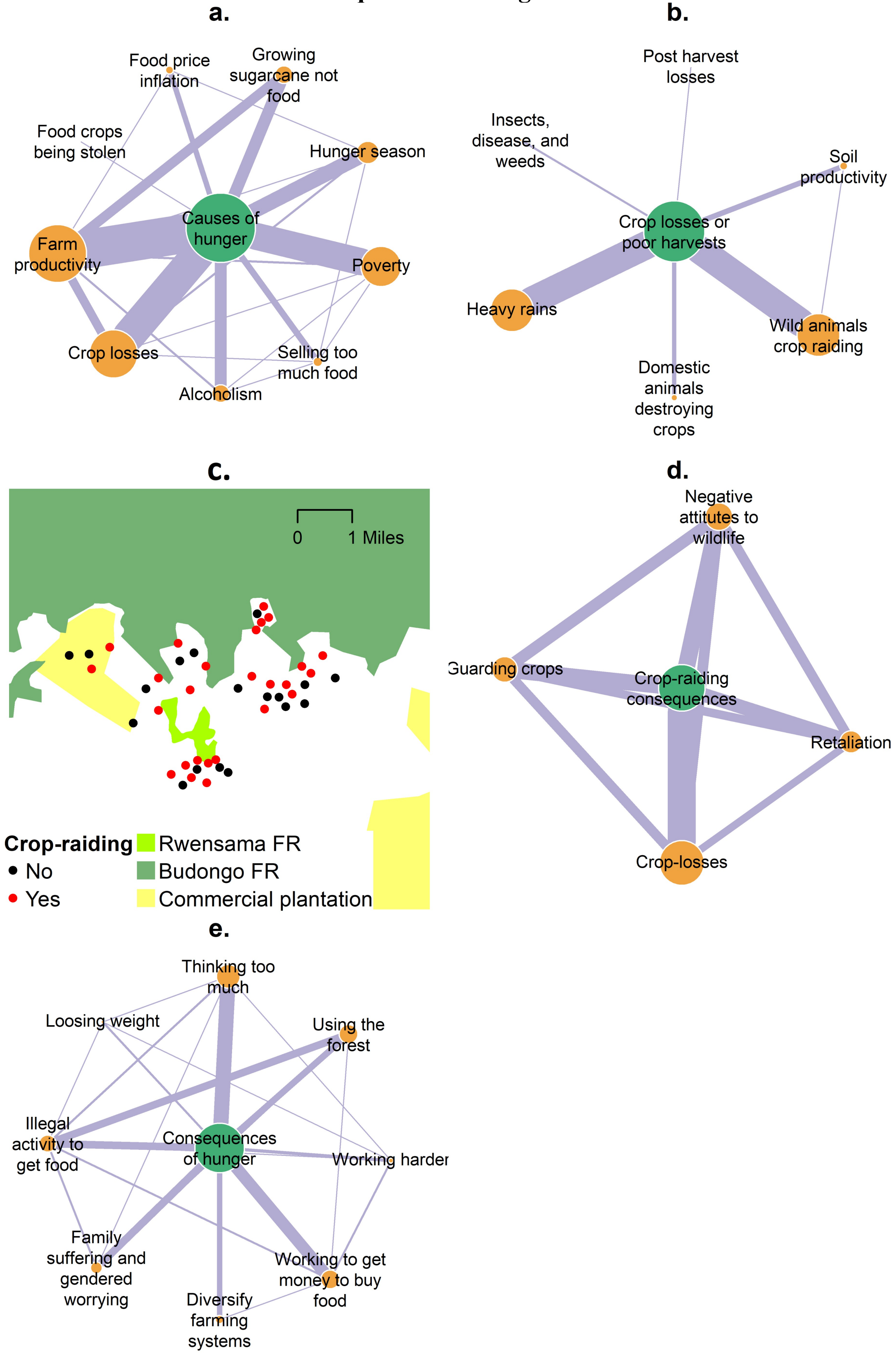


Figure 4. The width of the lines illustrates the number of interviews that reported connections between nodes but did not imply the direction of the causality of connections. The size of the node represents the number of

interviews mentioning the associated theme for that node. Panel a. illustrates the reported causes of hunger and inadequate food. Panel b. describes reported factors causing crop-losses. Panel c. shows the locations of interviews that report crop-raiding by wildlife (manually dislocated to retain anonymity). Panel d. illustrates the consequences of crop-raiding by wildlife, particularly among farmers bordering the Budongo and Rwensama Forest Reserves. Illustrates the reported consequences of experiencing hunger or inadequate food.

### S12: Forest use

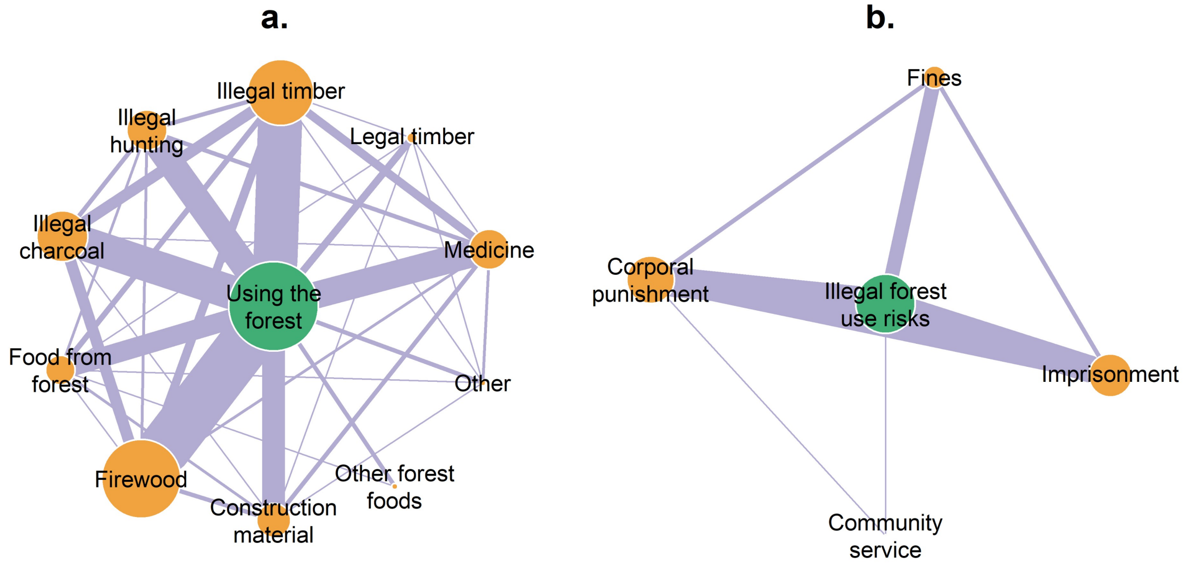


Figure 5. The width of the lines illustrates the number of interviews that reported connections between nodes but did not imply the direction of the causality of connections. The size of the node represents the number of interviews mentioning the associated theme for that node. Panel a. illustrates the reported uses of the forest. Panel b. describes the risks associated with illegal activities in the forest.
